# 3D single cell scale anatomical map of sex-dependent variability of the rat intrinsic cardiac nervous system

**DOI:** 10.1101/2020.07.29.227538

**Authors:** Clara Leung, Shaina Robbins, Alison Moss, Maci Heal, Mahyar Osanlouy, Richard Christie, Navid Farahani, Corey Monteith, Jin Chen, Peter Hunter, Susan Tappan, Rajanikanth Vadigepalli, Zixi (Jack) Cheng, James S. Schwaber

## Abstract

We developed and analyzed a single cell scale anatomical map of the rat intrinsic cardiac nervous system (ICNS) across 4 male and 3 female hearts. We find the ICNS has a reliable structural organizational plan across individuals that may provide the foundation for further analyses of the ICNS in cardiac function and disease. The distribution of the ICNS was evaluated by 3D visualization and using data-driven clustering. The pattern, distribution and clustering of ICNS neurons across all male and female rat hearts is highly conserved, demonstrating a coherent organizational plan in which distinct clusters of neurons are consistently localized. Female hearts had fewer neurons, lower packing density and slightly reduced distribution, but with the same localization. We registered the anatomical data from each heart to a geometric scaffold, normalizing their 3D coordinates for the standardization of common anatomical planes and providing the path where multiple experimental results and data types can be integrated and compared.

## Introduction

We recently published a comprehensive and cellular resolution 3D mapping of the rat intrinsic cardiac nervous system (ICNS)(Armour, 2008). The ICNS, regarded as the little brain of the heart, integrates multiple local sensory and autonomic efferent inputs and in turn regulates cardiac functions. We here extend the methods of approach to assay the structural consistency and variability of the rat ICNS within and between sexes. One of the most useful properties of other nervous systems – i.e. the brain – is that there is, to a first approximation, a regular and expected structural organizational plan across all brains of a species, and even to a useful extent across species. This allows data to be accumulated and compared across individuals and treatments. More recently, the development of digital brain atlases as quantitative data reference systems(Ding et al., 2016; Funka-Lea and Schwaber, 1994). To this end, we both evaluate ICNS for these uses and explore the use of the ICNS in a quantitative heart Scaffold.

The ICNS contains a variety of neurotransmitters and modulators that receive the inputs of both local afferent and extrinsic autonomic (parasympathetic and sympathetic) nerves, performing very complicated integration in controlling cardiac functions(Achanta et al., 2020; Ardell and Armour, 2016; Moss et al., 2020). We previously identified multiple clusters of ICNS neurons on the epicardium of the superior surface of the heart and on the posterior left atrium (Achanta et al., 2020; Ai et al., 2007; Cheng et al., 1999; Lin et al., 2008; Pauza et al., 2000; Richardson et al., 2003). While characterization of the ICNS shows that topology as well as neurotransmission to and from the heart is significantly conserved among species, little work has been done to examine the variability in the ICNS structure between individuals(Hopkins and Armour, 1984; Janes et al., 1986; Kawashima, 2005; Kuntz, 1930; Mizeres, 1963; Randall et al., 1972; Saccomanno, 1943). Additionally, while significant differences have been reported in the autonomic control of the cardiovascular system between males and females on a clinic level, possible differences in anatomical organization are not well understood(Fu and Ogoh, 2019; Kerkhof and Miller, 2018).

Although the study on sections and whole-mounts of the tissues has tremendously advanced our understanding on ICNS distribution and extrinsic and intrinsic nerve innervation, recent advances in imaging techniques have further improved the capability of visualizing the ICNS within its 3D anatomical framework. Exciting advances in the tissue clearing techniques have managed to visualize the ICNS and the autonomic axons in the whole heart(Chung et al., 2013). Following tissue clearing, immunohistochemistry and various microscopic techniques can be applied to map ICNS and extrinsic and intrinsic cardiac nerves in the whole heart(Rajendran et al., 2019). ICNS and extrinsic and intrinsic cardiac nerves may be imaged in high resolution with light sheet microscopy, as demonstrated on the murine heart(Ding et al., 2018). However, the penetration of light-sheet microscopy is limited to a few hundred micrometer thickness and not at a cellular resolution. To overcome these problems, we recently established a novel technical workflow that uses Knife Edge Scanning Microscope (KESM) in combination with the Tissue Mapper software we enhanced for this purpose that allows for comprehensive high resolution quantification of ICNS neurons and their qualitative visualization in the 3D spatial context of whole rat hearts(Achanta et al., 2020). Here, we built upon this proof-of-principle technique to explore the organization of the ICNS with respect to the anatomical features and uncovered the extent and distribution of ICNS neurons to compare and contrast the organizational scheme across individuals and sexes.

## Results

### Comprehensive 3D ICNS mapping to evaluate the variability of ICNS across individuals and between sexes

The comprehensive mapping of single ICNS neurons and their distribution were qualitatively and quantitatively analyzed for consistency and variability within and across the sexes. In this we apply the data acquisition pipeline developed in our recent paper (Achanta et al, 2020) that employs a Knife Edge Scanning Microscope (KESM) for high-resolution image acquisition and 3D ICNS mapping with the Tissue Mapper software we developed for this purpose; we were able to qualitatively and quantitatively visualize and examine the distribution of ICNS neurons in 3D reconstructed hearts as demonstrated in our previous study(Achanta et al., 2020) (Figure 1A). We applied the novel approach to perform comprehensive mapping of single cardiac neurons across four male and three female Fischer 344 (F344) rat hearts (Figure 1B, Video S1). From the overall distribution of ICNS neurons, the Partitioning Around Medoids (PAM) algorithm was used to assess the clusters present based on packing density of neurons throughout the ICNS. This analysis also resulted in the identification of neuronal clusters that were used as a guide to compare the anatomical organization of neuronal clusters across individual hearts (Figure 1B, Video S1, Video S2). We further extended our comparative analysis for two male ICNS by registering the 3D mapping data onto a mathematical representation of the heart known as a scaffold, which normalizes the 3D coordinates into a standardized framework of anatomical structures.

**Figure 1:**
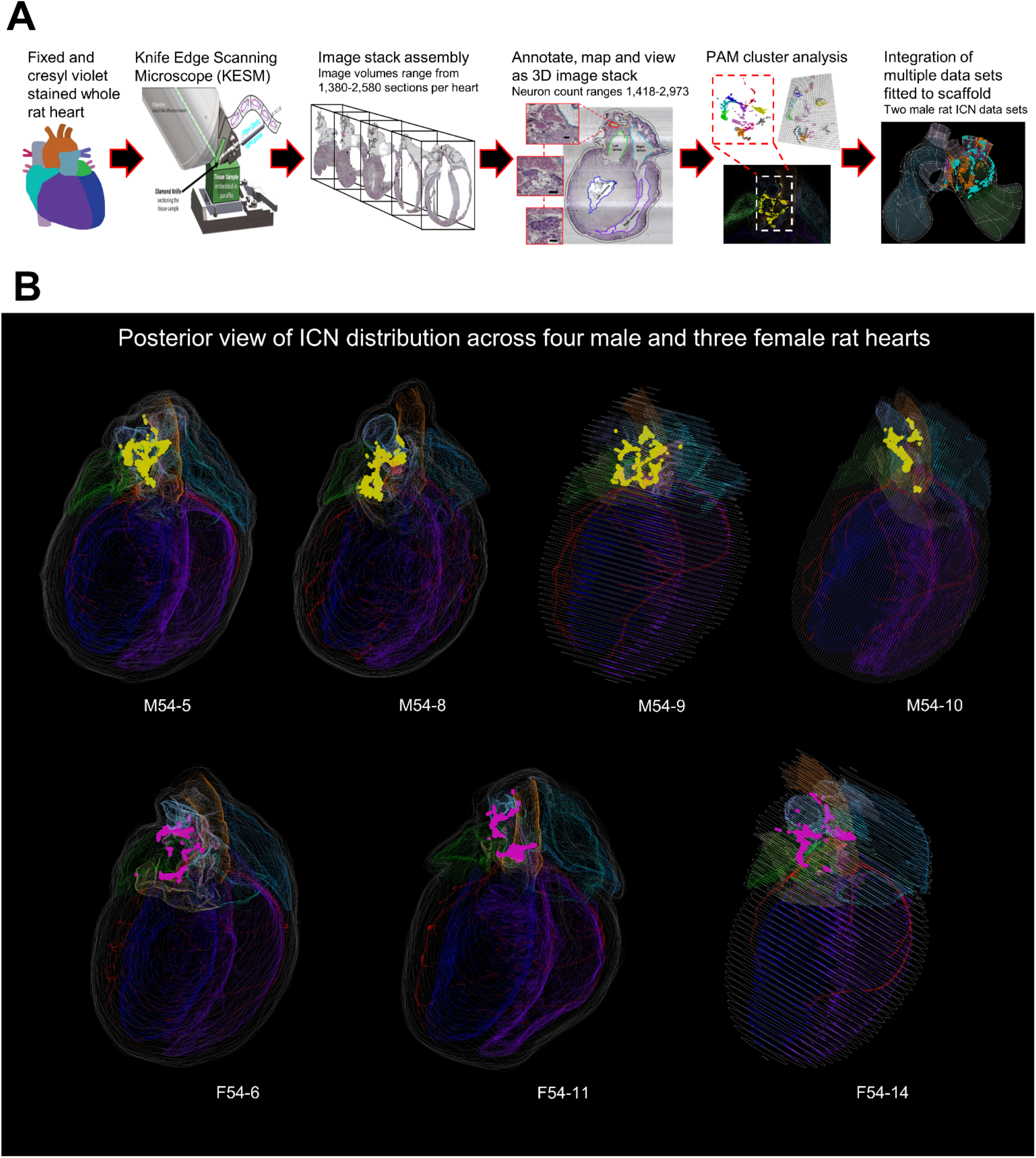
Spatially-tracked anatomical and molecular map of rat ICNS to enable comparison of variability within and across sexes. **(A)** Schematic representation of the workflow to acquire male and female rat ICNS maps, analysis of the data using cluster analysis of distinct groups of neurons, and alignment of datasets into a common coordinate framework. **(B)** Posterior whole heart views of four male and three female rat hearts. Yellow: males; Pink: females.

### The neuron clusters and distribution follow similar spatial pattern with variable density

A broad visual inspection of the ICNS neurons alone show they are not each randomly and individually distinct; they all form a limited number of clusters that are distributed three-dimensionally in a fundamentally similar fashion. At the same time there is variability between clusters in their shape, numbers, density, and specific distribution, somewhat reminiscent of the variability, for example, in the human brain’s cortex (Figure 2). In the male rat hearts (Figure 2A), large clusters of neurons were seen to be even distributed throughout the ICNS(Achanta et al., 2020), although some variations were observed (Figure 2A). Meanwhile, female ICNS consistently show a smaller number of large, distinct clusters of cardiac neurons, with lower neuronal packing density and with a few neurons scattered in between (Figure 2B). One male heart, M54-10, appears as an outlier due to having histological damage that lost part of the pulmonary veins, which contain clusters in all the other hearts. On average, female rat hearts presented with 1581 single ICNS neurons per heart, respectively (Figure 2C). This is a low estimate, as histological issues were present in some sections from female hearts preventing mapping. Without one of the male hearts (M54-10, 1722 mapped neurons; Figure 2C), the other three male rats were consistent with an average number of 2845 neurons, ranging from 2676-2973. Given that in these immature rats males are larger, the larger size may be consistent with greater numbers in males.

**Figure 2:**
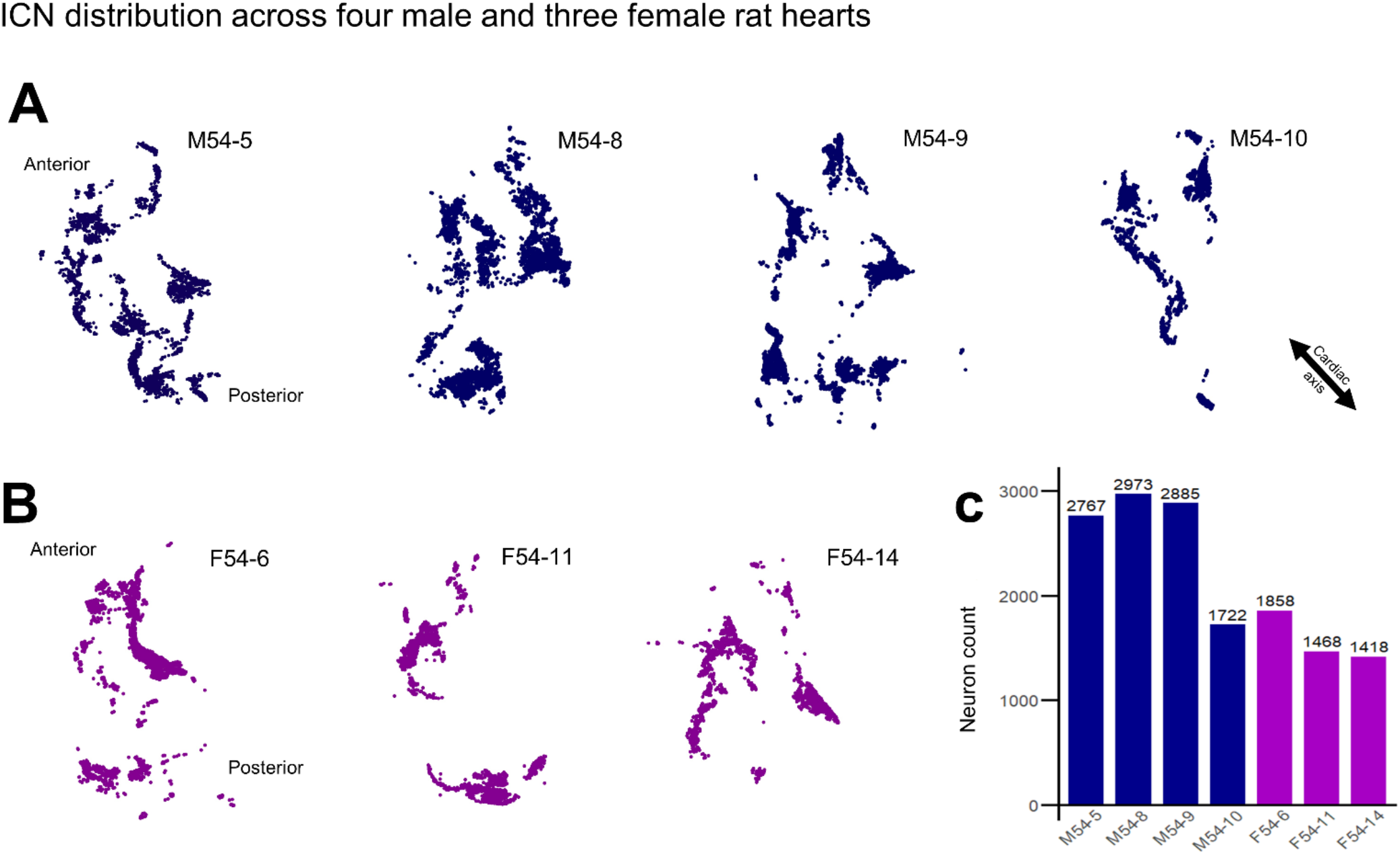
Quantity and distribution of neurons within the rat ICNS across individuals and sexes. **(A,B)** The anatomical map of rat ICNS across four males (A) and three females (B). Each dot represents a single neuron. The anatomical orientation corresponding to the whole heart is indicated. Blue: males; Purple: females. **(C)** Quantitative analysis of the total number of neurons mapped and annotated within seven rat hearts.

### ICNS neurons distribute around the same cardiac anatomical regions in male and female rats

Across all male and female rat hearts, the exact shapes and locations of the ICNS neuron clusters are somewhat variable, but these clusters are not randomly distributed around the heart but rather restricted, in all hearts, to the same cardiac anatomical Regions of Interest (ROI) on the base of the heart and the posterior surface of the left atrium (Video S2). We define the base of the heart as its superior or rostral border, including the hilum where the major great vessels access the heart. No neurons were identified in the ventricles. In order to more precisely compare and contrast the distribution of ICNS neurons between hearts, we used a quantitative and unbiased algorithm to identify different neuronal clusters. The partitioning around medoids (PAM) algorithm identified clusters of neurons based on packing density where the number of clusters was determined both mathematically and with the aid of visual inspection(Kaufman and Rousseeuw, 1987) (Figure 3A, Figure S1). Identification of clusters in each heart allowed us to more accurately identify and compare the distribution of neurons with respect to their surrounding specific anatomical features. When applied towards further examination of the neuron distribution in male rat heart M54-8, we observe clusters on the base of the heart and also a continuous band of cardiac neurons spanning the posterior or dorsal aspect of the left atrium that extends into a c-shaped ring that terminates at the base of the left atrioventricular sulcus (Figure 3A,C). These left atrium-associated neurons are also seen in histology of the ICNS (Figure 3B). In sum, multiple clusters were on the posterior left atrium and also bordered the neighboring coronary sinus and right pulmonary artery (Figure 3B). Using Neurolucida Explorer, we associated neuron clusters with these different anatomical features (Figure 3C). Five of the twelve PAM-identified clusters are associated with the posterior left atrium and the respective ICNS neurons have been highlighted to contrast to other clusters that were closer to other cardiac anatomy (Figure 3D). Through guided use of the PAM-identified clusters and by orienting the ICNS with respect to these features of cardiac anatomy in the remaining male and female hearts, we consistently observed neurons to follow the same patterns of distribution as denoted by the blue and purple colorations (Figure 3E-J).

**Figure 3:**
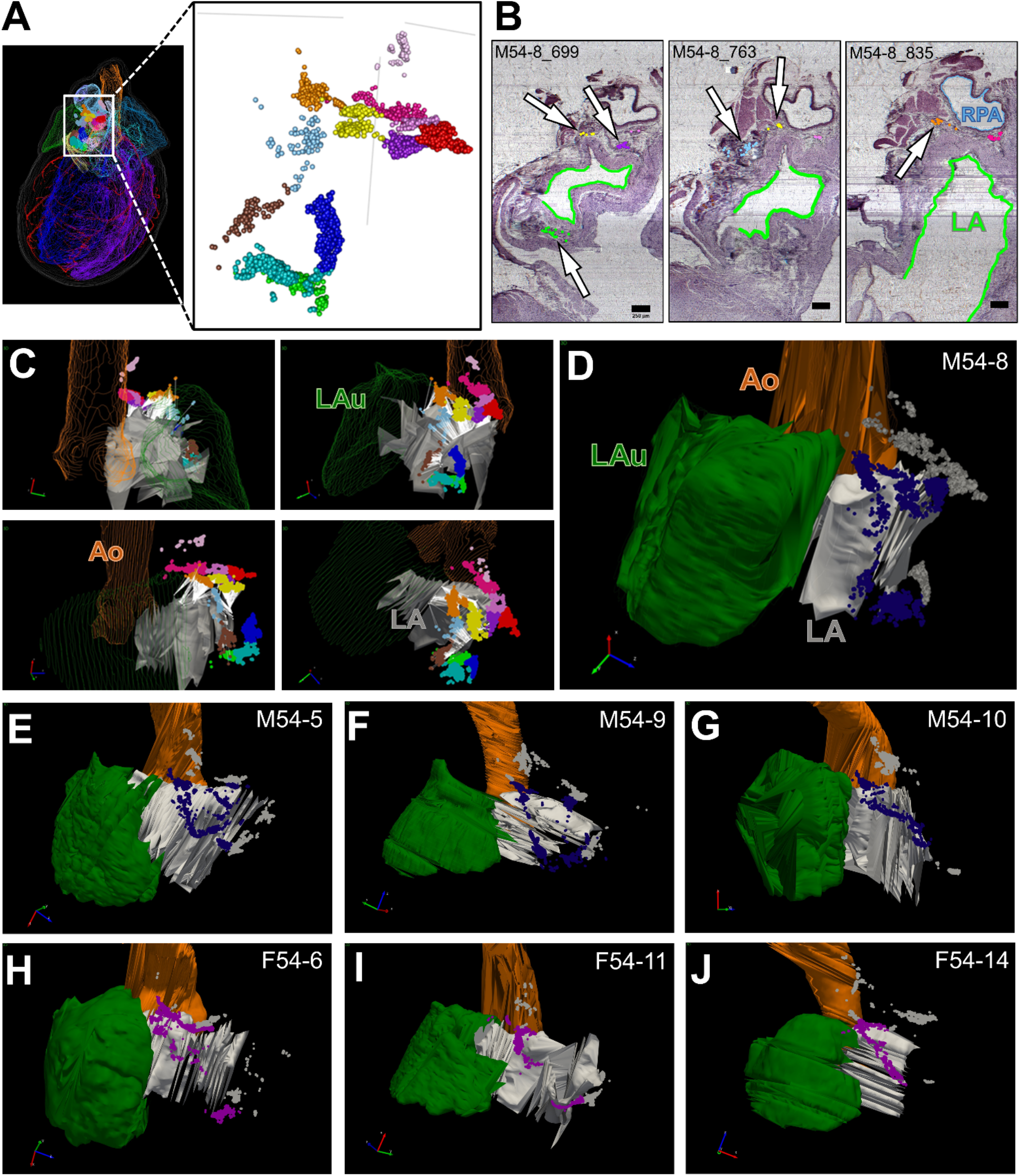
ICNS neurons are consistently localized around the left atrium. **(A)** Partitioning around medoids (PAM) clustering analysis in a male rat heart delimited 12 groups of ICNS neurons found at the base of the heart, visualized here with each color representing a distinct cluster of cells. (B) Histological sections throughout various levels of the image stack corresponding to the male rat heart shown in panel (A) were evaluated to identify which clusters were distributed around the left atrium. (C) In addition to the histological context, clusters near the left atrium were visualized in their 3D spatial context using Neurolucida Explorer as an independent analysis. **(D)** The PAM-identified clusters were re-colorized as dark blue and grey to represent ICNS neurons proximal and distal to the left atrium, respectively. **(E-J)** 3D spatial evaluation for the remaining male (E-G) and female (H-J) rat hearts identified the neurons that are proximal and distal to the left atrium. The neurons proximal to the left atrium are re-colorized as dark blue (male) and purple (female), whereas the neurons distal to the left atrium are visualized in grey. Scale bar: 250 μm.

### Data-driven cluster analysis of ICNS neurons on the base of the heart across male and female rat hearts also shows preferential distribution of neurons around regions of interest

In addition to the left atrium, we further studied the regional distribution of the ICNS and the degree of similarity between two male rat hearts, M54-5 and M54-8, using the PAM-identified neuronal clusters as a guide. We see similar regional distribution patterns at the hilum of the pulmonary veins entering the left atrium, the root of the superior vena cava and right atrium junction, the left atrioventricular sulcus (groove) as denoted by the coronary sinus, and the anterior interatrial sulcus of the left and right atria. In both male hearts, there were consistently distinct groups of neurons observed to form the characteristic c-shaped ring around the hilum of the pulmonary veins and left atrium (Figure 4A). Although fewer in number, ICNS neurons were consistently seen to be sparsely distributed along the root of the superior vena cava entry with the right atrium. These neuronal groups are also seen to wrap around the superomedial curvature of the right pulmonary artery (Figure 4B). Lastly, ICNS neurons were seen to cluster along the left atrioventricular sulcus (groove) around the coronary sinus, however the quantity of neurons present within that region was more variable between the two male hearts (Figure 4C). In addition to the ICNS neurons commonly observed within those three anatomical regions, a large number of neurons were also frequently seen to populate the anterior interatrial sulcus. Across both of the male rat hearts M54-5 and M54-8, ICNS neurons at the base of the heart distributed along the epicardial surface of the left and right atria as either one large group of cells or as multiple clusters (Figure 5A,B). By examining both hearts from a left superior angle of the posterior aspect, we can see that ICNS neurons within this area are situated directly beneath the right pulmonary artery (Figure 5C,D). Furthermore, the histological sections provide additional spatial context of these ICNS neurons in relation to other anatomical features within this region (Figure 5E,F). Additional cluster analysis of ICNS neurons in these four anatomical regions of interest additionally extending to male rat hearts M54-9 and M54-10 and to female rat hearts F54-6, F54-11, and F54-14 all demonstrated similar distributions within those regions (Figure 5G). While there is a fine grain variation between individuals and across sexes, our analysis indicates that in the above ROI, on the posterior left atria and in specific locations on the base of the heart, there is consistency in the principle organization structure guiding the regional distribution of ICNS neurons.

**Figure 4:**
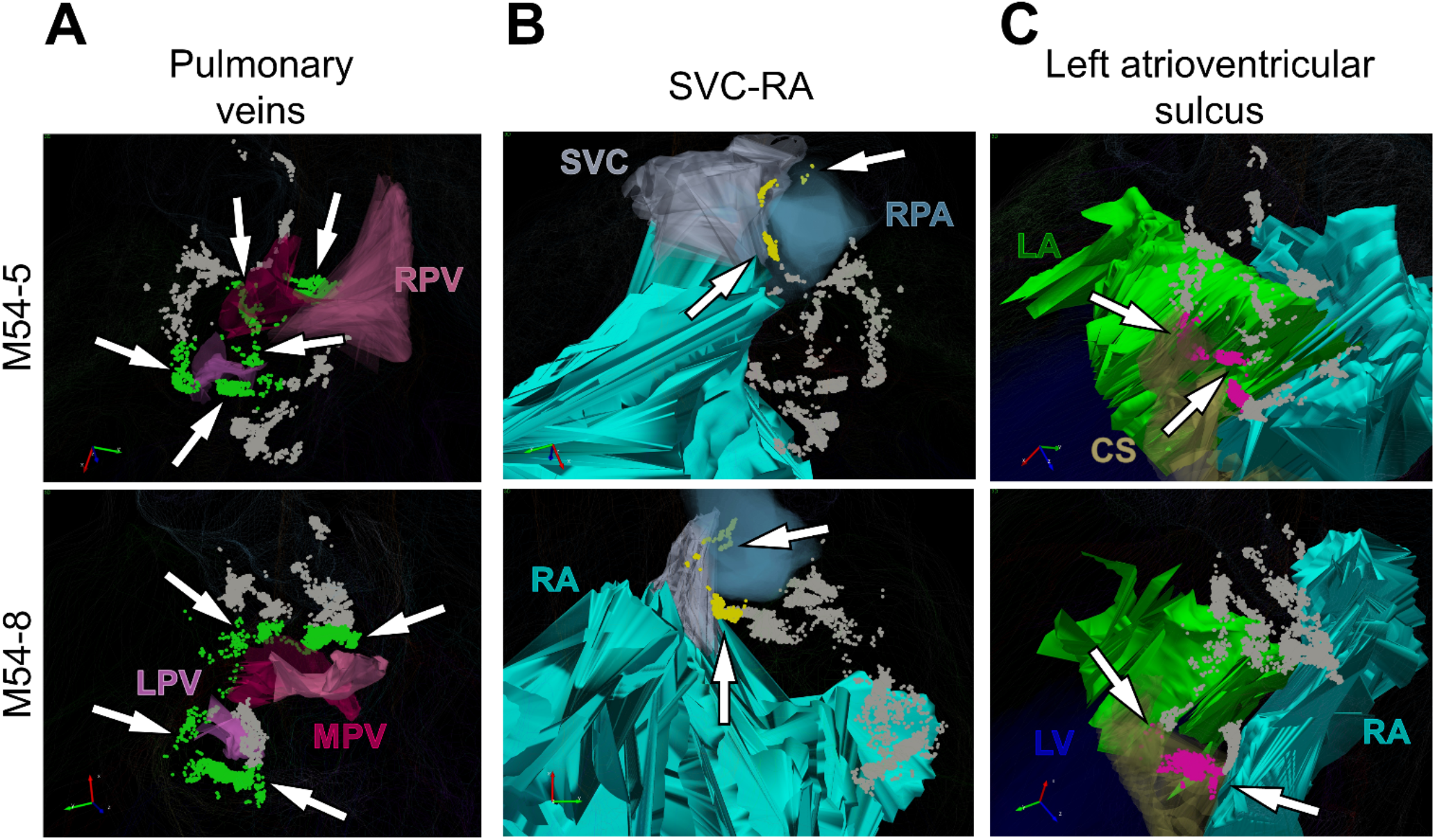
Data-driven analysis of ICNS demonstrates preferential organization of ICNS neurons in select cardiac anatomical regions. **(A-C)** ICNS neurons from two male rat hearts that organize around the pulmonary veins (PV) (A), the root of the superior vena cava and right atrium (SVC-RA) (B), and the atrioventricular sulcus (C). **(A)** ICNS neurons encircling the pulmonary veins highlighted in green. **(B)** ICNS neurons around the root of the superior vena cava and right atrium (SVC-RA) are highlighted in yellow. **(C)** Clusters of ICNS neurons present at the left atrioventricular sulcus near the coronary sinus are highlighted in pink. In (A-C), all neurons not emphasized as being associated with the particular region of interest are shown in grey.

**Figure 5:**
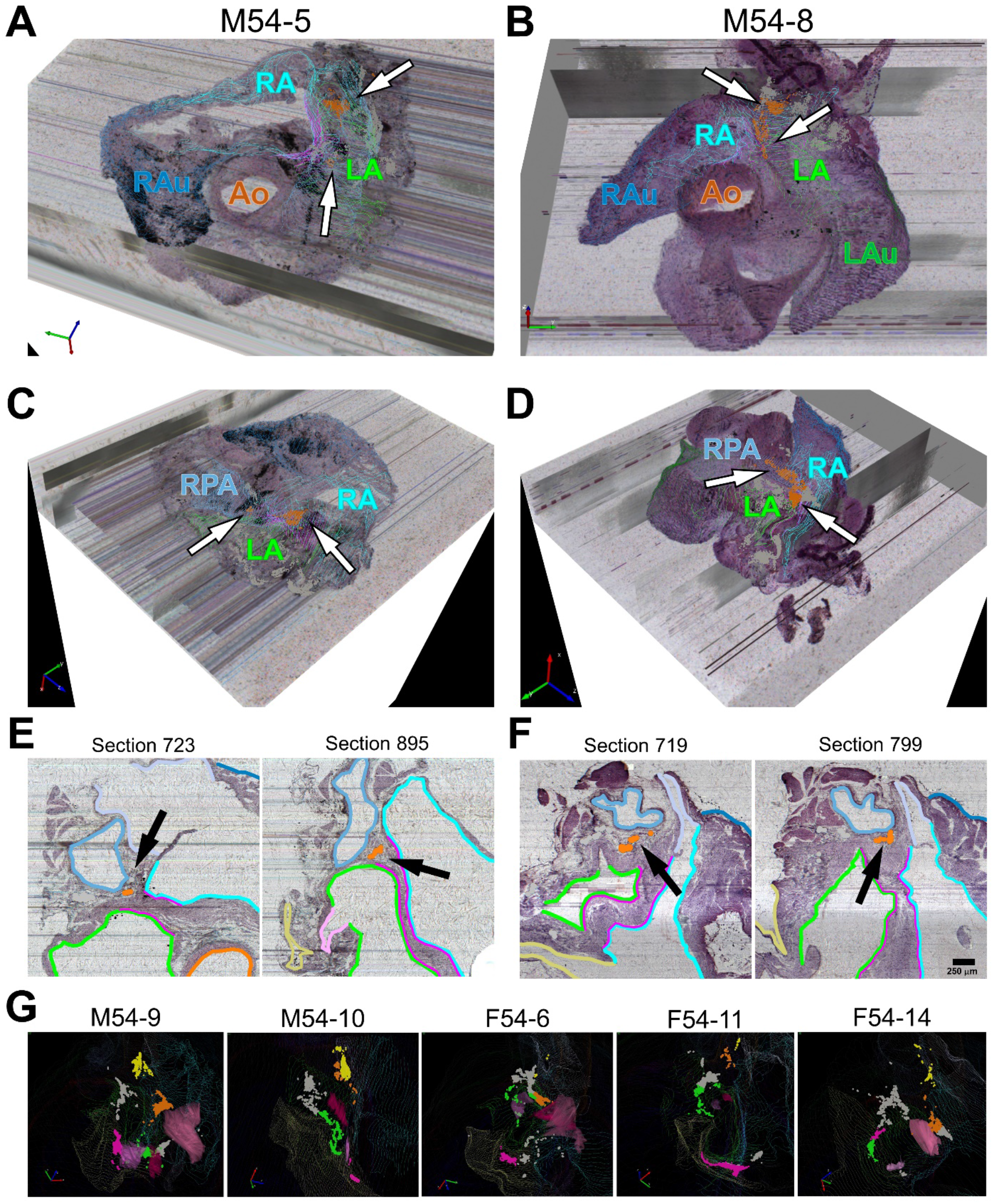
ICNS neurons are localized along the interatrial sulcus. **(A-G)** Partial projections and histology for M54-5 (A,C,E) and M54-8 (B,D,F). **(A,B)** A superior view of partial projections of a selective thickness of heart tissue visualized as 3D image volumes for a pair of male rat hearts. Overlaying contours of the left and right atria onto the ICNS highlights a large number of ICNS neurons distributed along the anterior interatrial sulcus. **(C,D)** A left superior view of the posterior aspect of the partial projections illustrates the distribution of ICNS neurons along the anterior interatrial sulcus that are obscured by the right pulmonary artery. **(E,F)** Cardiac histology shows the location of specific clusters in relation to the left and right atria as well as the right pulmonary artery. Scale bar: 250 μm. (G) Such a pattern of localization to these cardiac anatomical regions was observed across all the male and female rat hearts. The colors green, yellow, and pink correspond to ICNS neurons around the PV, the root of the SVC-RA, and the left atrioventricular sulcus, respectively (From Figure 4).

### Integration of two datasets into a generic heart scaffold identifies regions of dense overlap

We continued the assessment of ICNS similarity between male rat hearts M54-5 and M54-8 by fitting the 3D tracing data and mapped ICNS onto individual heart scaffolds (Figure S3). A scaffold is a mathematical approach to represent the standard shape of an organ, in this case the heart, through a 3D material coordinate system. Within this coordinate framework, many aspects of the heart can be represented including the musculature, vasculature, and the spatial distribution of the ICNS. The advantage for fitting the 3D mapping data into the scaffold is the ability to compensate for distortions to the image volume and corresponding tracings by accurately and mathematically representing the native structure of the heart while maintaining the structural anatomy and spatial position of the ICNS (Figure 6A, Figure S3, Video S3).

**Figure 6:**
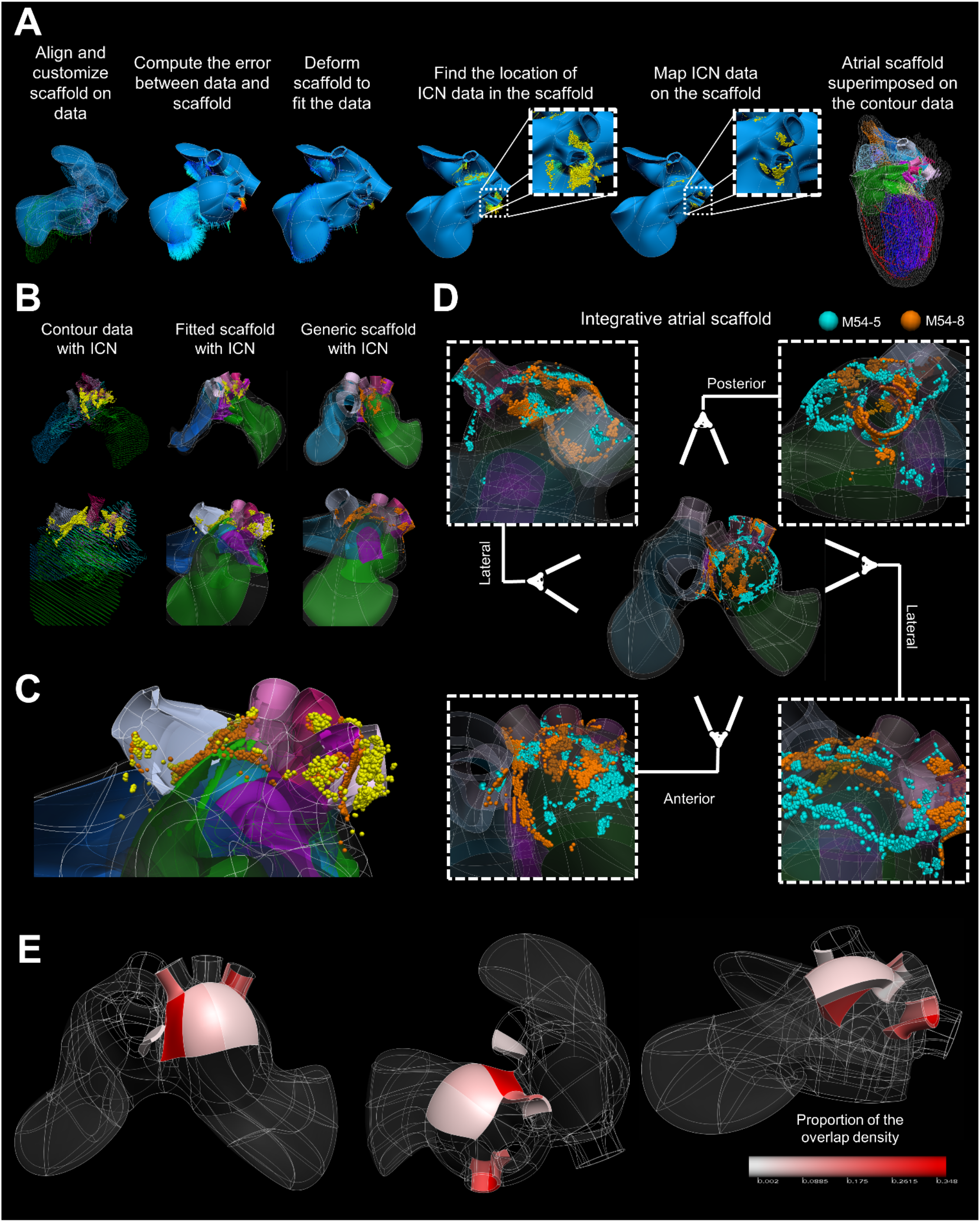
Mapping individual rat ICNS and cardiac anatomy onto a generalized 3D scaffold for comparison across animals. **(A)** Workflow for registering individual ICNS into a 3D scaffold, fitting the scaffold to the individual cardiac anatomical features, and projection of individual cells on the scaffold to embed the ICNS in its elements. **(B)** An overview of the fitting and mapping processes from the original 3D anatomical segmented data of one of the male rat hearts viewed from two different angles. The original contour data, fitted scaffold and generic scaffold are shown. **(C)** The heart is visualized from a left superior angle to appreciate how the neuron locations in the original data are projected onto the generic scaffold. Yellow spheres represent positions of the original ICNS locations prior to transforming the data onto the fitted scaffold; orange spheres represent the new ICNS locations embedded in the scaffold. **(D)** Integration of two data sets onto one generic scaffold. **(E)** Looking at the same regions of interest in figures 4 and 5, varying colors of pink are used to indicate the proportion of neuronal overlap between the two male datasets from panel (D).

The 3D volumetric data of the base of M54-8 anatomically segmented heart with spatially recorded ICNS neuron locations are aligned with a heart scaffold (Figure 6B). The scaffold is then deformed to minimize errors and improve fitting accuracy to the 3D anatomically segmented data. Once the scaffold is in the same coordinate space as the original data, ICNS neurons are mapped onto the scaffold as element material coordinates (Figure 6B, Figure S4). Looking more closely at the left superior aspect of the heart, we can see how the positions of the neurons from the original 3D anatomical segmented data are projected onto the nearest surface of the scaffold for an accurate fitting which can then be integrated into the generic scaffold of the heart (Figure 6C, Figure S4). By integrating the anatomical and mapped ICNS data from two male rat hearts into one common coordinate space, we are able to visualize Regions Of Interest at the base of the heart with a high proportion of overlapping ICNS neurons (Figure 6D, Video S3). Furthermore, analysis of the overlapping regions of the ICNS across animals suggests results consistent with the similarity of ICNS distribution in the four ROI, around the left atrium, the left and right pulmonary veins, the superior vena cava, and the interatrial sulcus (Figure 6E, Video S3). Integration of these datasets onto the scaffold allows for quantitative comparison between and across multiple species despite variation in the topographic organization of the ICNS.

### Quantitative analysis of male and female ICNS demonstrate a similar organization between individuals and across sexes

Although there is a difference in the number of neurons observed between male and female rat hearts, the distribution pattern of ICNS neurons are present in the same ROI at the base of the heart and posterior left atrium. To describe the similarities and differences in the organization of the ICNS between sexes we compared the ICNS of male rat heart M54-8 with that of female rat heart F54-6. From the whole-heart perspective of both the male and female rat hearts, we see once again that ICNS neurons are distributed on the posterior left atrium (Figure 3, 7A,B). By orienting the ICNS of both the male and female rat hearts in a 2D packing density plot, aided with identification of high density clusters through PAM analysis, we see that the greatest difference presents in the relative numbers of neurons in various locations, rather than their overall distribution. (Figure 7C,D, Figure S1,S2). In the male rat heart, there is a uniform packing density of ICNS neurons throughout the superior-inferior extent with no sharp peaks present in any particular group of cells. In contrast, the female rat heart exhibited a higher proportion of ICNS neurons at the superior aspect of the heart (Figure 7C,D).

**Figure 7:**
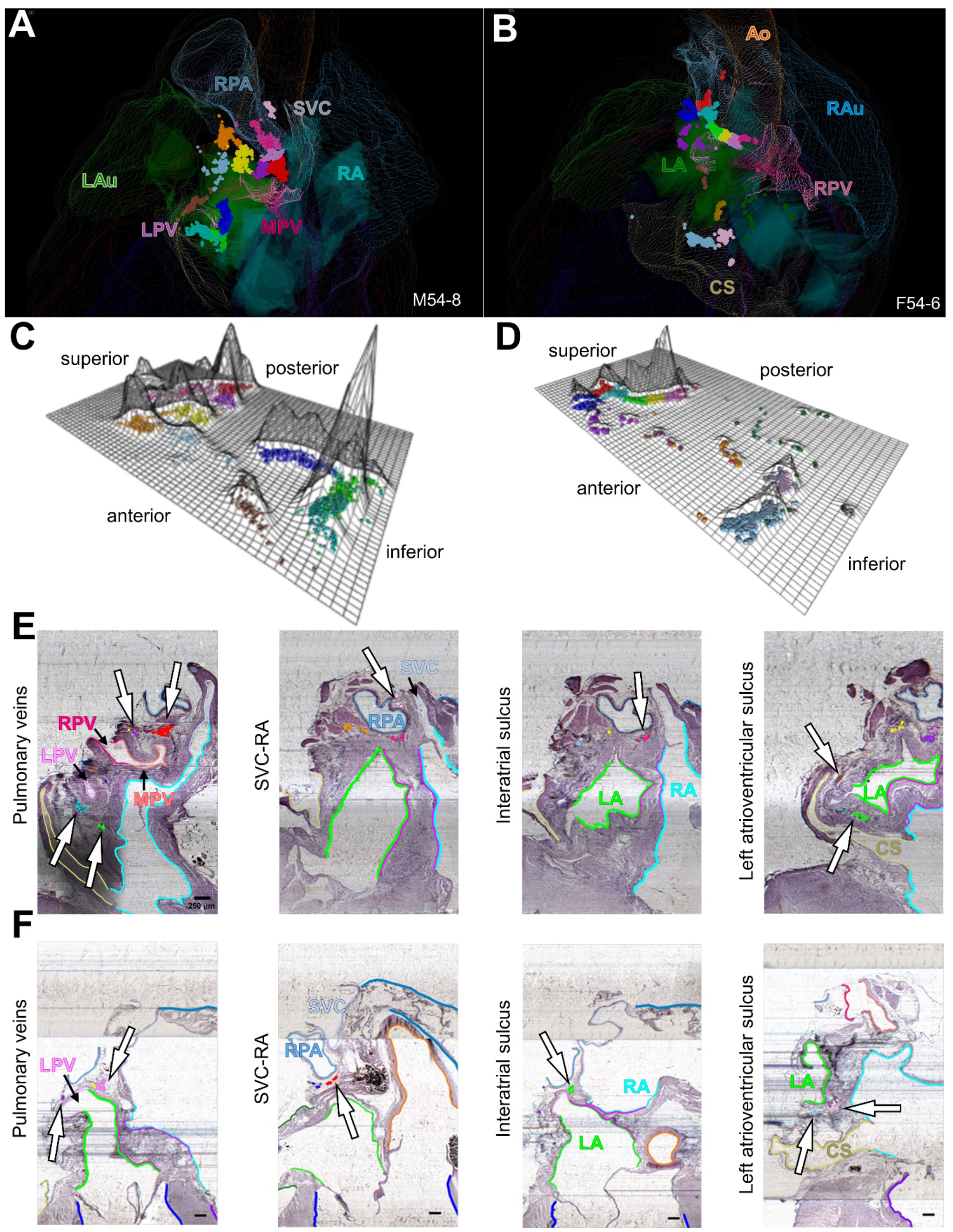
Comparison of male and female ICNS. **(A,B)** Whole heart posterior view of PAM clustered neurons of male (A) and female (B) ICNS. **(C,D)** Flatmap projections derived from PCA plots are used to show the spatial data and packing density of the neurons in male (C) and female (D) ICNS colored for PAM-identified clusters. **(E,F)** Histological tissue sections containing PAM colored neurons in relation to the four regions of interest are shown in a 2D context to support the 3D visual data for male (E) and female (F) ICNS. Scale bar: 250 μm.

Examination of the histology at the hilum, interatrial sulcus, and atrioventricular sulcus shows corresponding PAM-identified neuronal clusters within those regions of interest for both the male and female rat hearts, supporting our observation that there is a preferential organization in the regional distribution of ICNS neurons (Figure 7E,F, Figure S5,S6, Video S4,S5). While the male and female rat hearts exhibit differences in the packing densities of ICNS neurons, the localization pattern and extent of the neurons illustrates consistent organization between individuals and across the sexes.

## Discussion

In this study, we applied our recently established novel approach on the comprehensive mapping of the ICNS that uses a KESM for high-resolution image acquisition, the Tissue Mapper software for single neuron mapping and cardiac annotation, and PAM algorithm for cluster identification and density packing. Together, this workflow allowed us to qualitatively visualize and quantitatively determine the spatial distribution of ICNS neurons within and between sexes in 3D digitally reconstructed whole hearts(Achanta et al., 2020). We found that the pattern, distribution, and clustering of ICNS neurons are highly conserved in both males and females, though variability in terms of the exact location, shape, and number of ICNS neuron clusters is present. Compared to males, our present data strongly suggests that the female ICNS comprises significantly fewer neurons, although the localization and distribution of ICNS neurons were largely overlapping at the base of the heart at matching Regions Of Interest (ROI) in both males and females. To further develop our novel ICNS mapping approach, we have created generic heart scaffolds that are mathematical representations of the native organ and subsequently fitted the ICNS of two male rat hearts onto a single integrative scaffold such that the distribution and variability of cardiac neurons can be compared in a common heart model. Together, our data demonstrates a robust organizational plan of the ICNS that provides an anatomical foundation for future comparative, integrative, and functional studies of brain-heart connectome, molecular phenotype, and chemical coding features of the ICNS.

While there was variability in the ICNS distribution between individuals and sexes, we found that neurons were consistently organized around four distinct regions of interest: the hilum of the left atrium and the pulmonary veins, the root of the right atrium and superior vena cava junction, the left atrioventricular groove, and the anterior interatrial sulcus. As observed in multiple species, such an organization suggests a connection between anatomical features and their functional relationship. In large animals (rabbit, dog, pigs), functional studies showed the two large ganglionic plexuses (GP) of the right atrial ganglionated plexus (RAGP) and the middle pulmonary vein and caudal vena cava (PVCV) may regulate sinoatrial (SA) and atrioventricular (AV) functions to some extent(Allen et al., 2018; Arora et al., 2003; Cardinal et al., 2009; Hardwick et al., 2014; Nakamura et al., 2016; Pauza et al., 2014, 2000; Petraitiene et al., 2014; Saburkina et al., 2014; Singh et al., 2013; Steele et al., 1994; Xi et al., 1991). In rats, two large clusters of ICNS have been reported at similar locations as in the rabbits and pigs, which also regulate the SA heart rate and AV conduction(Ai et al., 2007; Cheng et al., 2004, 1999; Cheng and Powley, 2000; Hoard et al., 2008; Li et al., 2014, 2010; Pauza et al., 2000; Richardson et al., 2003; Rysevaite et al., 2011b, 2011a; Sampaio et al., 2003)\ Our work, mapping out all ICNS neurons, expands upon the concept of ganglionic plexuses with dedicated functional targets by suggesting that while the existence of these GPs may be consistent across individuals and species, the organization of each GP at the single neuron level is highly variable. This organization necessitates annotating the neurons in the ICNS based on how they participate in control of the end organ, not based purely on their location and implies that while the neurons may not be found in the exact same location in every individual, they will have a similar relationship with the potential downstream end-organ targets.

It has been well known that there are significant differences in neuro-hormonal regulation of the cardiovascular system in males and females(Kerkhof and Miller, 2018). Sex differences in baroreflex sensitivity and prevalence of cardiovascular disease in pre-menopausal women may be related to the ICNS differences we observe. It will be interesting to consider future studies that examine the relationship between neurons and packing density of neurons with their functionality in a diseased state. The pattern of ICNS distribution in the female rat hearts appeared more variable than in the male hearts and were present in far fewer numbers and at a lower density. Despite this, we consistently found ICNS neurons located in the four major regions of interest in both males and females, indicating that neurons in the ICNS of both sexes congregate in similar anatomical regions. Although there is variability in the density and distribution of the neurons in each of these regions, their presence suggests that it is not control of the functionality of the ICNS that differs between males and females but rather the strength of that control. The number of neurons necessary to participate in the baseline functional circuits may differ between sexes, and difference in density may explain some of the sex differences observed at a functional level, especially when moving away from the baseline healthy state to a state of injury or disease.

In performing this first study using this data acquisition pipeline, we encountered several histological issues that can affect the quality of the data. One of them is apparent in M54-10 where a piece of the hilum and associated ICNS were cut off during removal. This emphasizes the care that must be taken with post-mortem removal of the organs. In addition, particularly in the female hearts, variable staining quality in some cases obscured ICNS clusters in such a way as to make them incompletely mapped. While this may have some effect on the lower number of neurons seen in the ICNS, there are several regions that were successfully mapped in full that still showed a lower density than seen in the males. We have now worked out staining methods that will resolve these issues in future studies. Improvement of tissue preparation may lead to more similarity including locations, shapes, sizes, packing density and number of neurons. For example, the atrial tissue and the major vessels are soft and thus the shape of the left and right atria as well as the major vessels were not identical in size and shape. Future normalization of atria after fixation will reveal more similarity of ICNS distribution within and across sexes. However, given the variation of ICNS neuron distribution from the whole-mounts of many carefully prepared male animals, variability between individuals is expected (Li et al., 2014; Rysevaite et al., 2011b, 2011a). Additionally, based on learnings in the present application of PAM, we now are prepared for future studies to further adapt the analysis to consider anatomical features surrounding neuronal clusters in order to partition them without the aid of visual inspection.

From the integration of the ICNS of two male rat hearts into one integrated scaffold, we have demonstrated a replicable and reusable framework for the incorporation of additional information on the functional, molecular, and phenotypic aspects of the ICNS, which will provide the foundation for a brain-heart connectome atlas. Projection of multiple datasets onto a common scaffold allows for integration of pre-established findings throughout the known literature to be integrated into a common coordinate framework, providing a path to more efficiently build upon findings obtained in previous studies. Placing the ROIs identified in our study into the scaffold as a standard reference system and further integrating anatomical, electrophysiological, and more data types into a unified representation of the heart allows all of the findings to be studied as a single system. Eventually, the integrative scaffold can present a complete cardiac-brain atlas with all anatomical, physiological, and molecular information for cardiac control. This integration of multiple relevant data types in a common framework provides a natural substrate to enable modeling and simulation studies to be done in a more systematic structure, as is being done through the O^2^S^2^PARC platform (Neufeld et al., 2020). Whether aging and diseases may remodel ICNS differently is an important issue in development of targeted therapeutics. Through modeling and simulation of the ICNS we can further dissect the variability in the organization of the ICNS across individuals and between sexes and how it is remodeled in pathological conditions. These developments will inform and accelerate the progress towards neuromodulation therapies and further enable the development of treatments tailored to each individual.

## Limitations of the Study

- The anatomical annotations and representations may contain interobserver variability due to the manual segmentation process. The outlines and borders of the anatomic structures depend on subjective interpretation to some extent, which is unavoidable in a manual annotation process.
- Unbiased clustering of single neuron positions through PAM analysis was aimed at the whole heart and not specific cardiac substructures. The PAM clusters were interpreted visually for proximity and adjacency to cardiac anatomical features.
- Histological issues such as efficiency of staining can affect the quality of the data due to incomplete mapping in a few sections. We attempted to mitigate such issues by comparing across sequential sections to mark individual neurons.

## Data and Code Availability

The authors declare that all the data supporting the findings of this study are available within the article and its supplemental information files or from the corresponding author upon reasonable request.

All sample acquisition images and annotations pertaining to 3D spatial location are publicly available in the sparc.science (RRID:SCR_017041) repository with the digital object identifiers 10.26275/g6wb-qc7q and 10.26275/wcje-hxib.

A dataset containing high-resolution figures, supplemental figures, tables, and movies as well as the TissueMapper XML annotations are available at 10.26275/jvmm-45ly.

PAM algorithm for clustering and analysis are available as part of the R software (RRID:SCR_003388) in the package *cluster.*

Scaffold-Maker is an open-source Python library package developed in-house at the ABI to mathematically represent the generic atrial topology of the heart and is available at: https://github.com/ABI-Software/scaffoldmaker/.

Fitting of the scaffold with the 3D mapping data was performed with the OpenCMISS-Zinc platform developed at the ABI and is available at: http://ooencmiss.org/.

Development of the workflow management system was achieved using the MAPCIient package and is available at: https://github.com/MusculoskeletalAtlasProject/mapclient.

## Acknowledgements

Financial support for this work was provided by the National Institutes of Health (NIH) under the Stimulating Peripheral Activity to Relieve Conditions (SPARC) program grant OT2 OD023848 (PI: Kalyanam Shivkumar, UCLA) subaward to J.S.S., R.V. and Z.C.; National Heart Lung and Blood Institute (NHLBI) grant U01HL133360 to J.S.S. and R.V.; NIH grant OT3OD025349 to P.H.; and, NHLBI grant R15 1R15HL137143-01A1 and National Institute of Neurological Disorders and Stroke (NINDS) grant U01NS113867 to Z.C.

## Author contributions

J.S.S., R.V., Z.C., and S.T. conceived and designed the original workflow and corresponding experiments. M.H. and S.T. provided technical expertise on Tissue Mapper usage and facilitated in the coordination of data curation with MAP-CORE. M.H. created movies for animated visualization of the data. C.L. performed the 3D mapping and cardiac feature annotation. J.C. assisted in sample preparation. N.F. and C.M. performed the staining and KESM imaging. S.R. performed the PAM data analysis and comparisons for within and across sex differences, and figure assembly. A.M. wrote the code for the PAM analysis. R.C. developed the generic heart scaffold in which the 3D mapping data was fitted onto. M.O. performed computational registration of the 3D mapping data onto the heart scaffold. C.L., S.R., and A.M assembled the figures and wrote the manuscript with feedback from all the authors. P.H. supervised MAP-CORE efforts for mapping data onto the heart scaffold. Z.C. supervised the cardiac feature annotation. R.V. supervised the data analysis. J.S.S. supervised the overall study.

## Supplemental Information

**Figure S1:**
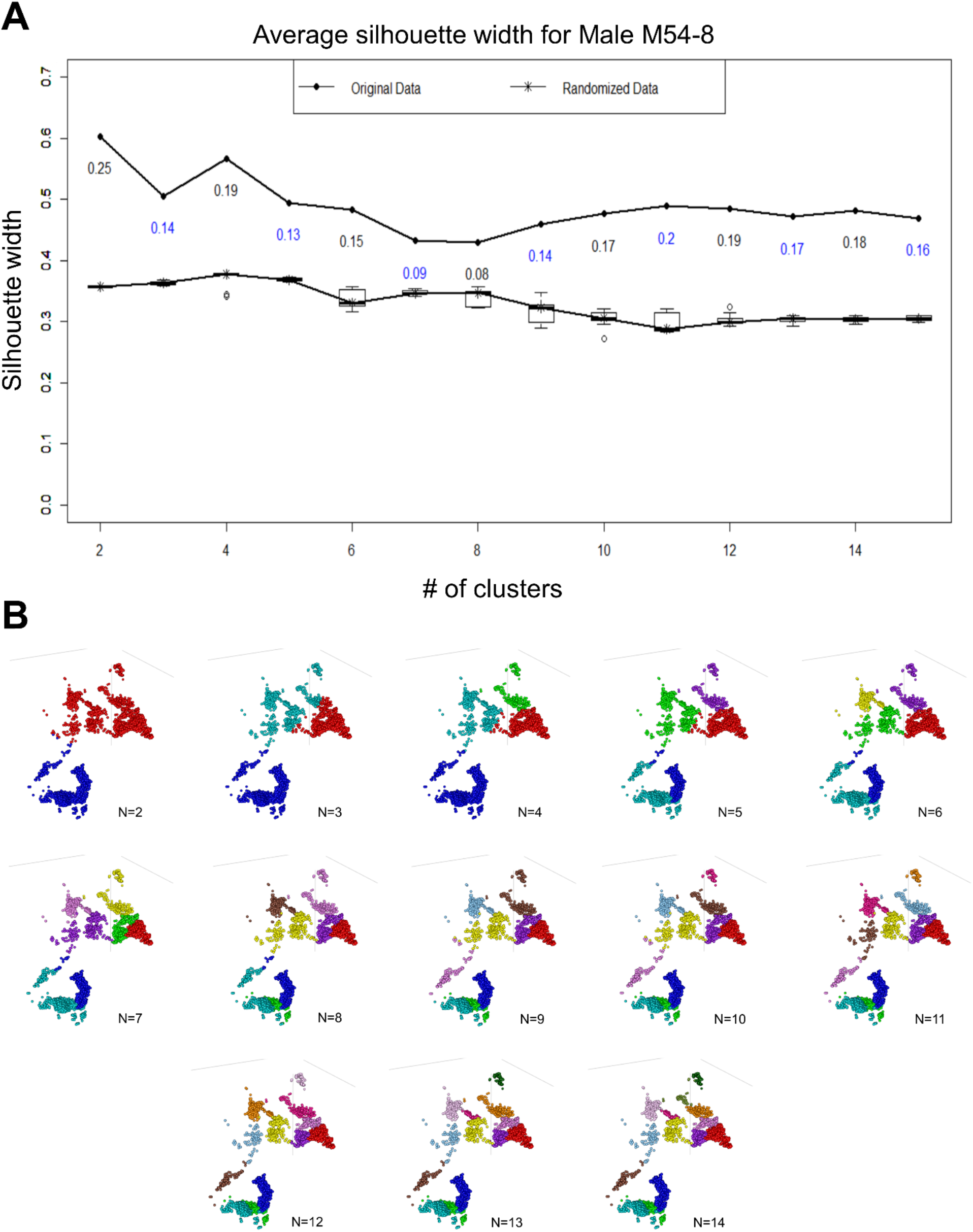
PAM analysis in male ICNS, related to Figure 3A; Figure 7C. Partitioning around medoids (PAM) analysis was performed on the ICNS coordinates for each cluster number, resulting in a silhouette coefficient describing how well the clustering fit the data. The coordinates were additionally randomized and subjected to the same analysis ten times to compare the silhouette width of randomized data to the original coordinates. **(A)** Graph comparing the silhouette width for both original and randomized data against the cluster number for male heart M54-8. The numbers below the line of original data in the plot shows the difference between the silhouette width of the original data and the mean of the randomized data. **(B)** For all possible cluster numbers, ICNS representations are colored based on the PAM assigned clustering.

**Figure S2:**
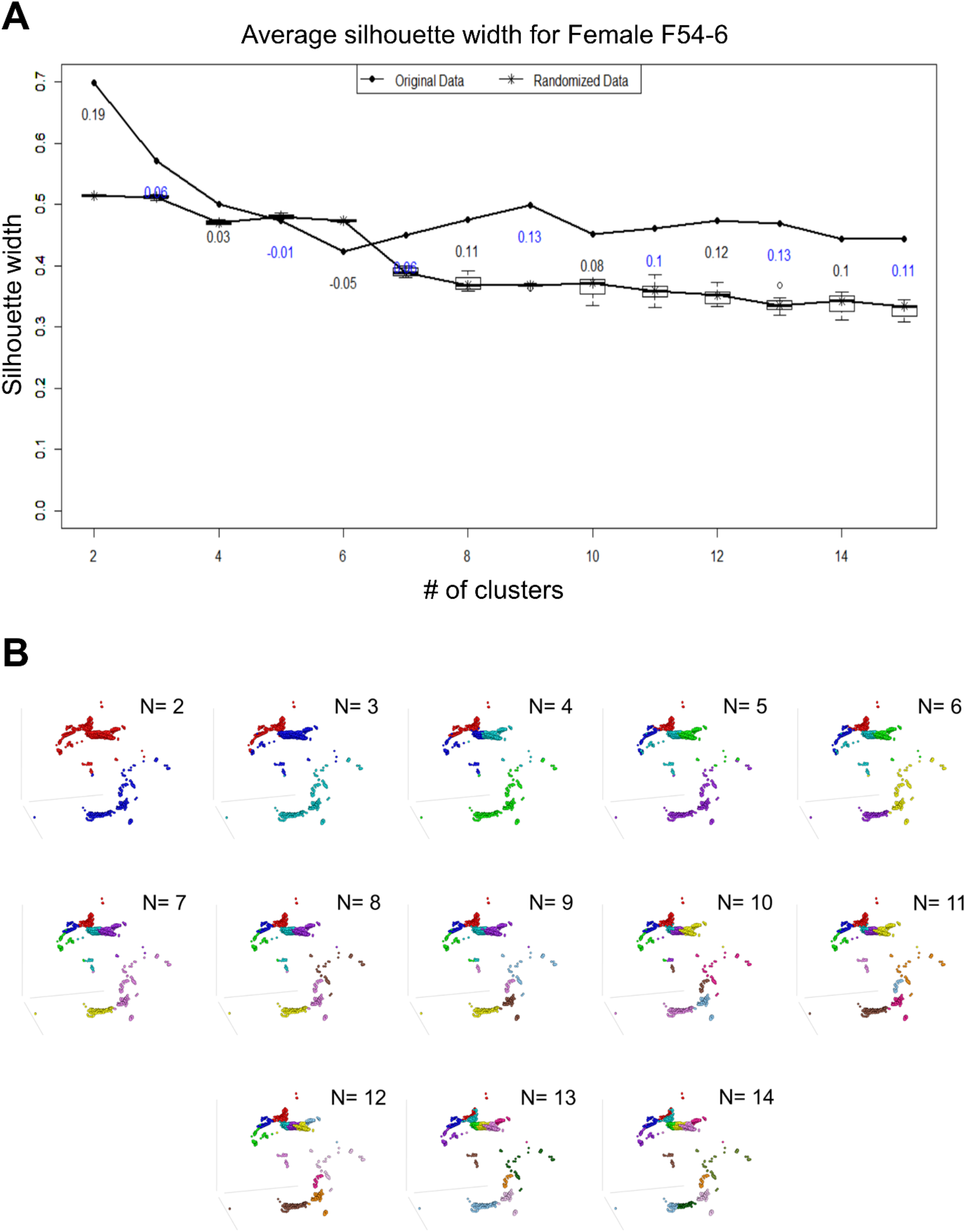
PAM analysis in female ICNS, related to Figure 7D. Partitioning around medoids (PAM) analysis was performed on the ICNS coordinates for each cluster number, resulting in a silhouette coefficient describing how well the clustering fit the data. The coordinates were additionally randomized and subjected to the same analysis ten times to compare the silhouette width of randomized data to the original coordinates. **(A)** Graph comparing the silhouette width for both original and randomized data against the cluster number for female heart F54-6. The numbers below the line of original data in the plot shows the difference between the silhouette width of the original data and the mean of the randomized data. **(B)** For all possible cluster numbers, ICNS representations are colored based on the PAM assigned clustering.

**Figure S3:**
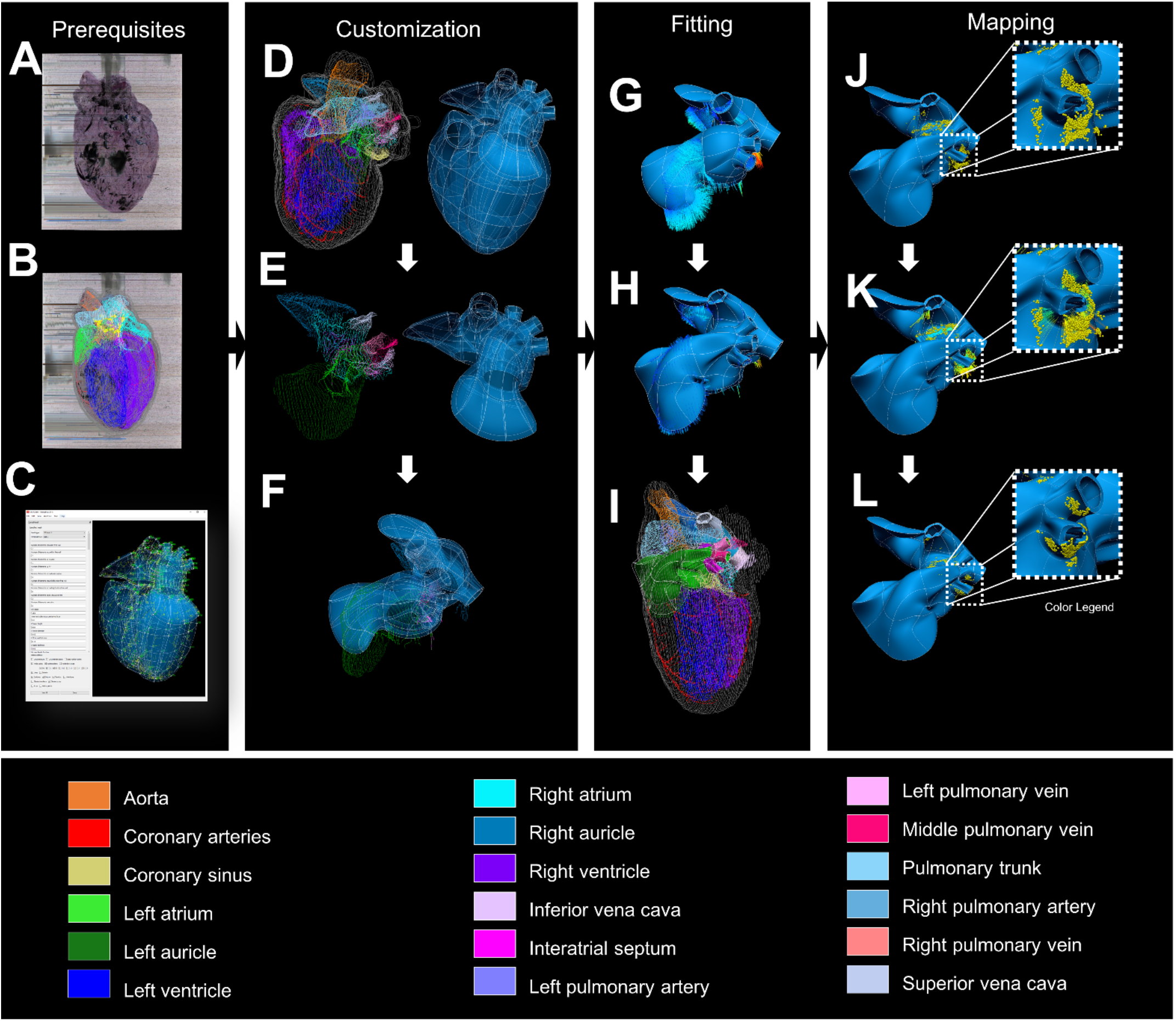
Scaffold fitting pipeline, related to Figure 6A, Video S3. **(A-C)** Three fundamental prerequisites are required for the mapping process. A volumetric image of the heart (A) is reconstructed to segment the regions into tracing contours (B). A corresponding scaffold is then generated from the Scaffold Maker software interface (C) to match the contour data. **(D-F)** Each heart must be customized to match the scaffold parameters to the data. The contoured heart data and the corresponding heart scaffold (D) are trimmed to include only the atria and auricles, pulmonary veins, superior vena cava, inferior vena cava, and interatrial septum (E). The scaffold is then aligned to the data through a rigid transformation process (translation, rotation, and scaling), and its parameters are tweaked empirically to provide the closest matching shape to the data (F). **(G-I)** the data must be fit to the scaffold and includes deformation of the scaffold to obtain an accurate shape of the data. Every point on the contour is projected onto the closest point on the surface of the scaffold (G) to find the sum of the Euclidean distances of these projections. This sum is then minimized through an iterative optimization process which deforms the scaffold to “fit” the data (H). The resulting fitted scaffold is superimposed on the original heart contour (I) for a further qualitative validation. **(J-L)** The ICNS data is then mapped onto the fitted scaffold. With the scaffold in the same coordinate system as the original data, the ICNS can be mapped onto the scaffold as element “material” coordinates. Some ICNS neurons are within the scaffold elements and others are wrapped around it (J). Those within preserve their exact spatial locations while the ones “hanging” around are projected to the nearest surface using a nearest search algorithm (K). This mapping results in embedding of ICNS into the scaffold and preserving their spatial distribution and cluster variation (L). Color legend shows the colors of every anatomically labelled region.

**Figure S4:**
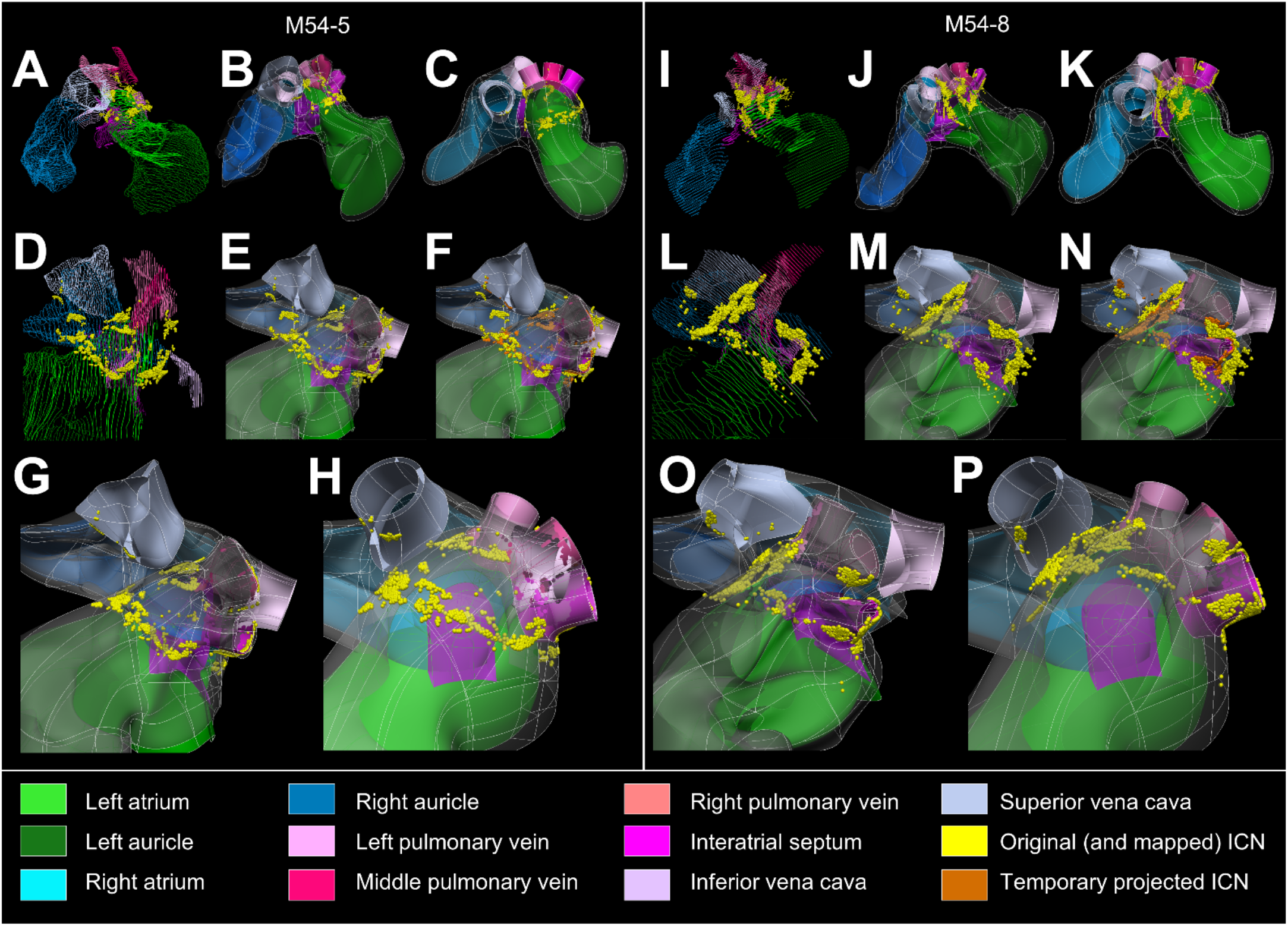
Mapping of ICNS from two male hearts onto the heart scaffold, related to Figure 6B,C, Video S3. Mapping results for the two male rat hearts: 54-5 (A-H) and 54-8 (I-P). **(A-C,I-K)** Big-picture representations of the original contour data (A, I), fitted scaffold (B,J), and generic scaffold (C,K) with ICNS neurons shown in yellow spheres. ICNS data is mapped into the element material coordinates for (B,J) and (C,K). **(D-H,L-P)** Close-up, superior views from the left side of the data and scaffold. Distribution of ICNS neurons above the left atrium and around the pulmonary veins (D,L) and ICNS neurons transformed onto the corresponding fitted scaffolds (E,M). Projection of ICNS onto the surface of the fitted scaffold (F,N) The [temporary] orange spheres represent the mapped locations of the original locations of the ICNS shown in yellow spheres. The exact location of any yellow sphere within the scaffold is preserved while for those yellow spheres located outside of the scaffold, an estimated location is found by projecting the ICNS neurons to the nearest surface plane. The resulting map of the ICNS data on the fitted (G,O) and generic scaffold (H,P). Note that the [temporary] orange color is changed back to yellow for consistency. The results show how this approach can provide the means of mapping and storing the spatial locations of the ICNS neurons within a quantifiable reference frame while keeping the pattern of ICNS distribution intact, and how this reference frame can provide means for comparing variation between different species.

**Figure S5:**
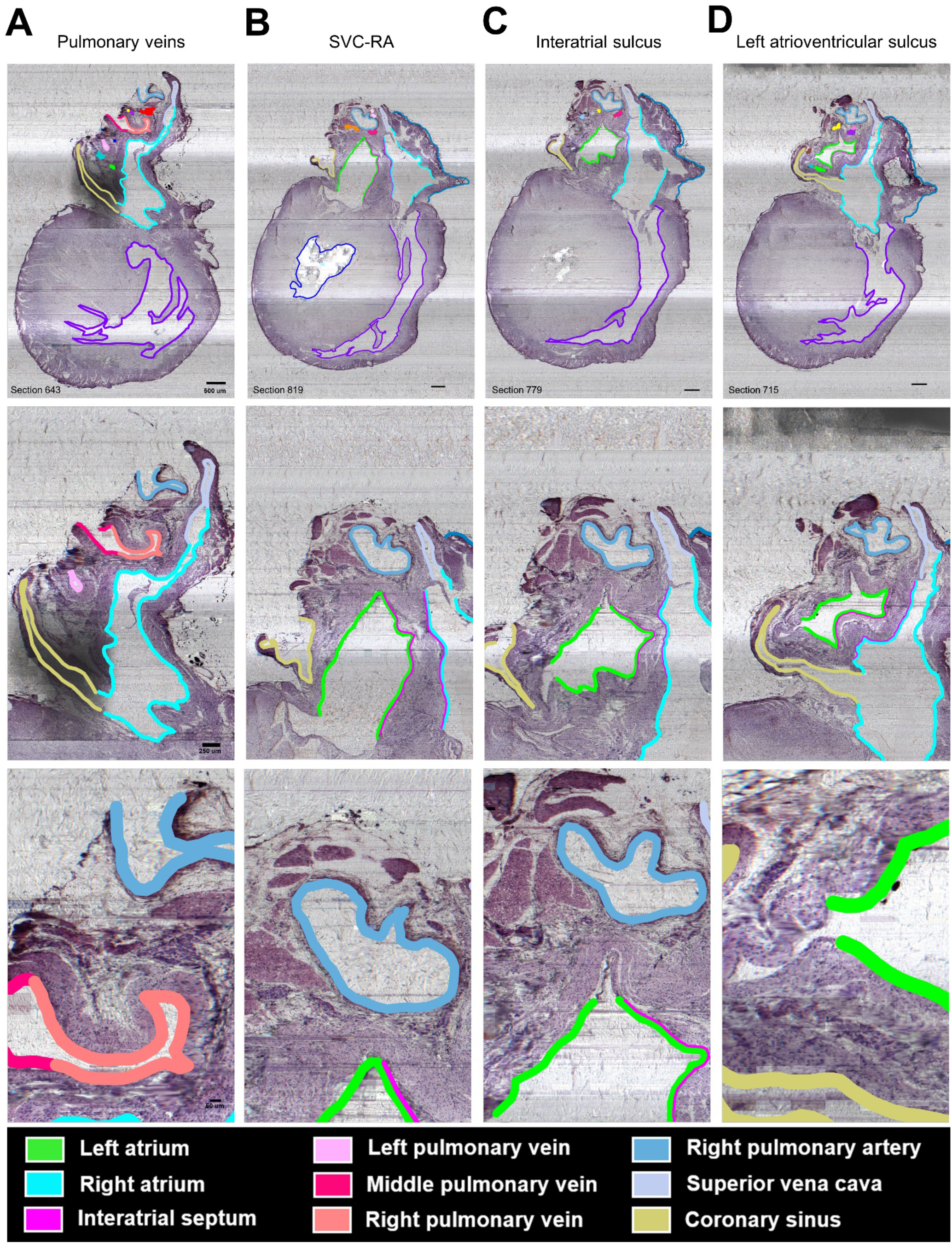
Histological sections of the male heart, related to Figure 3B, 5F, 7E, Video S4. **(A-D)** Histological sections from male heart M54-8 focusing on the pulmonary veins (A), superior root of the vena cava and the right atrium (B), interatrial sulcus (C), and left atrioventricular sulcus (D) at increased zoom levels from top to bottom.

**Figure S6:**
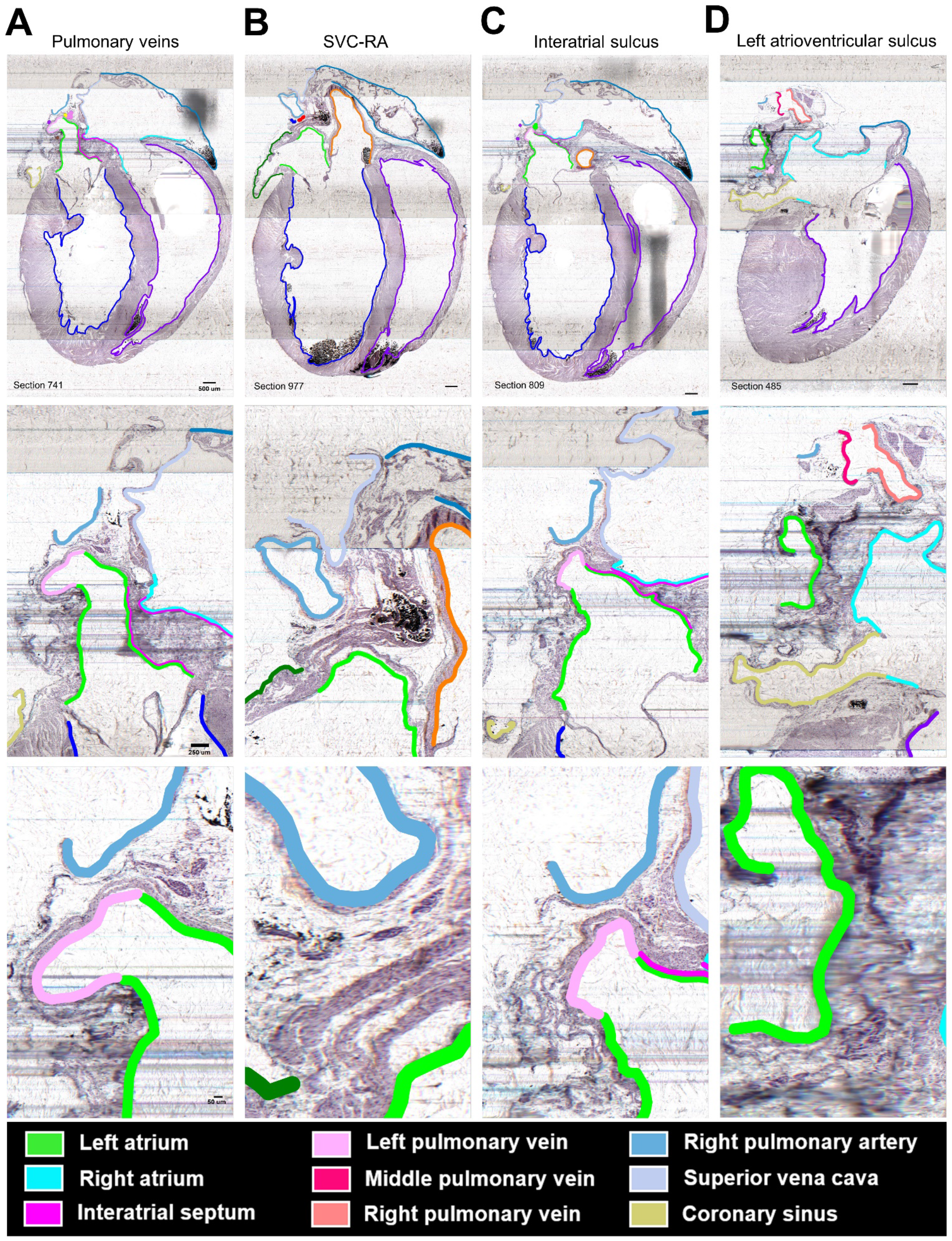
Histological sections of the female heart, related to Figure 7F, Video S5. **(A-D)** Histological sections from female heart F54-6 focusing on the pulmonary veins (A), superior root of the vena cava and the right atrium (B), interatrial sulcus (C), and left atrioventricular sulcus (D) at increased zoom levels from top to bottom.

**Table S1.** related to Figure 2: Details on the neuron count, number of sections in the image stack, and sectioning plane for each heart.

| Rat ID | Sex | Total neuron count | Average number of neurons (M vs. F) | Total number of sections in image stack | Total number of sections in image stack containing tissue | Sectioning plane |
| --- | --- | --- | --- | --- | --- | --- |
| M54-5 | Male | 2,676 | 2,564 | 1,400 | 1,210 | Parallel to long axis (coronal) |
| M54-8 | Male | 2,973 |  | 1,570 | 1,350 | Parallel to long axis (coronal) |
| M54-9 | Male | 2,885 |  | 2,580 | 2,580 | Parallel to short axis (transverse) |
| M54-10 | Male | 1,722 |  | 1,570 | 1,559 | Parallel to long axis (sagittal) |
| F54-6 | Female | 1,858 | 1,581 | 1,650 | 1,494 | Parallel to long axis (coronal) |
| F54-11 | Female | 1,468 |  | 1,380 | 1,344 | Parallel to long axis (coronal) |
| F54-14 | Female | 1,418 |  | 2,460 | 2,448 | Parallel to short axis (transverse) |

**Video S1, related to Figure 1B:**

Whole heart animation of four males (top) and three females (bottom) illustrating the anatomical maps of intrinsic cardiac nervous system.

**Video S2, related to Figure 1B, 7C,D:**

Comprehensive intrinsic cardiac nervous system mapping in the rat heart to enable comparison of variability within and across sexes.

**Video S3, related to Figure 6B-E, Figure S3, S4:**

Mapping individual rat intrinsic cardiac nervous system (ICNS) and cardiac anatomy onto a generalized 3D scaffold for comparison across animals.

**Video S4, related to Figure 3B, 5F, 7E, Figure S5:**

Flyby visualization of histological sections comprising the image stack for male heart M54-8 detailing the dimensions and size of the data obtained.

**Video S5, related to Figure 7F, Figure S6:**

Flyby visualization of histological sections comprising the image stack for female heart F54-6 detailing the dimensions and size of the data obtained.

### Transparent Methods

#### Animals and sample preparation

Four male and three female Fischer 344 (F344) rats four weeks of age were obtained from Envigo. Animals were anesthetized using 5% isoflurane. Once the animal was non-responsive to a contralateral toe pinch, the abdominal cavity was opened for subsequent perfusions and removal. For samples M54-5 and F54-6, the hearts were removed and fixed overnight in 4% paraformaldehyde before whole-mount diffusion staining was performed with Cresyl-Echt Violet (0.05g in 50mL dH2O and 150μL glacial acetic acid) for seven days to visualize intrinsic cardiac neurons. The protocol was updated and altered slightly for the remaining samples. The animal was perfused with phosphate buffered saline until exsanguinated via the ascending aorta at a pressure of 280 mmHg. Animal was then perfused at the same pressure with 200 mL 10% neutral buffered formalin. Hearts were dissected, further fixed overnight in 10% neutral buffered formalin, and whole-mount diffusion staining was performed with Cresyl-Echt Violet (0.05g in 50mL dH2O and 150μL glacial acetic acid) for fourteen days to visualize intrinsic cardiac neurons. The hearts were processed in a Sakura Tissue Tek VIP 3000 tissue processor, and then embedded in paraffin. A syringe was used to inject molten paraffin into chambers of the heart through great vessels in order to avoid air bubbles within the chambers, which can cause problems in KESM imaging. All procedures were performed in compliance with the National Institute of Health Guide for the Care and Use of Laboratory Animals. Protocols can be found at dx.doi.org/10.17504/protocols.io.bdz5i786.

#### Knife Edge Scanning Microscope (KESM) image acquisition, post-processing, and ICNS mapping in 3D reconstructed hearts with Tissue Mapper

The paraffin-embedded hearts were digitized with a Knife Edge Scanning Microscope (KESM) by sectioning heart tissue at a 5 μm thickness for each z-slice. The paraffin-embedded hearts were mounted onto a nano-precision XYZ stage that moved the sample along a diamond ultramicrotome knife coupled with a fiber-optic cable. The fiber-optic coupled microtome knife illuminated the edge of the knife where the tissue sample was sectioned. A custom-built objective with a 5mm field of view and a tube lens with magnification equivalent to 10x was focused on the beveled edged of the diamond knife. The tube lens was connected to a CMOS TDI line scan color sensor with a 16K pixel resolution RGB output with a 5μm x 5μm pixel size.

For each sample, the mounted paraffin-embedded hearts were moved by the robotic XYZ stage against the 5mm diamond knife to section the heart. As the heart was sectioned along the Y directionality, one continuous line or strip of image data at the X dimension was captured by the line scanning camera and generated a single image tile of 10,000 pixels where each pixel represented 0.5μm. After one strip of image data was captured, the stage then moved to an adjacent region of the heart and the process was repeated until the sample at that specific z-level was digitized. This whole process was repeated until the entire heart was sectioned.

The collected image data was post-processed with KESM software to denoise and normalize the background. The KESM generated image tiles from each XY location at every Z-level. The tiles were automatically aligned and stitched into 2D image planes, which represented one section of the heart. The pixels that did not contain the heart were cropped to remove excess image data. Each individual 2D image plane was then assembled into a 3D image volume with a 40:1 JPEG2000 compression (Biolucida Converter, MBF Bioscience, Williston). The image volumes contained a range of 1380-2580 sections per image volume that was dependent on the sectioning plane.

The image volumes were then annotated (Tissue Mapper, RRID:SCR_017321) to quantify and mark the location of intrinsic cardiac neurons and to annotate cardiac anatomy in the 3D image volumes. The annotation of cardiac anatomy was selected from a few structures that represented the chambers and major blood vessels found consistently in all hearts to simplify the comparative distribution of neurons between individuals. On sections where the major features were present (left and right atria, auricles, ventricles, aorta, pulmonary trunk and arteries, left, middle, and right pulmonary veins, superior vena cava, inferior vena cava, coronary sinus, and coronary arteries), the anatomy was traced with different colors for each representative feature in intervals across all the sections in which they were present. The surfaces contoured for the cardiac chambers and blood vessels were the endocardium and the tunica intima, respectively. The epicardium was contoured for the left and right auricles. The traced anatomy were visualized as 3D wireframe reconstructions in the Tissue Mapper 3D to compare the ICNS between individuals and across sexes. Detailed protocols can be found online at protocols.io as 10.17504/protocols.io.bdz5i786.

#### Intrinsic cardiac neuron mapping and quantification

ICNS neurons were mapped based on whether neurons were well-stained, if the eccentric nucleus was visible, if the cell body was unobscured by artifacts, and the cell size. ICNS neurons were identified by Nissl staining, morphology, and localization around specific cardiac anatomy and were mapped in intervals of four sections to prevent double counting. If the nucleus was not visible in the fourth interval section of mapping, then the section above and below was examined for a neuron with a visible nucleus that occupied the same position in the interval section of counting. Lastly, neuron somata were measured for a short and long axis dimension of at least 13 μm x 23 μm as the size criteria for cell mapping(Cheng et al., 1997; Pauza et al., 2002). All ICNS neurons were mapped and quantified by following the histological criteria for identifying neurons and the results are summarized in Table S1. Protocols can be found online at protocols.io as 10.17504/protocols.io.bdz5i786.

#### Cluster analysis using Partitioning Around Medoids (PAM)

PAM analysis was performed using the “cluster” package in R, using Euclidean as the distance metric. The PAM algorithm is similar to the k-means algorithm that works to break the dataset into groups and attempts to minimize the distance from each point to its designated group (Kaufman and Rousseeuw, 1987). Unlike k-means, k-medoids or PAM chooses a data point as a center. It minimizes a sum of pairwise dissimilarities instead of a sum of squared Euclidean distances. It makes the locally optimal choice at each stage with the intent of finding a global optimum point. In layman’s terms, the centers, or medoids of the clusters are chosen so as to minimize the distance between that center and surrounding points, essentially these centers will be in the most dense areas. Each point is then assigned to the medoid “group” by finding the minimum distance between that point and each medoid. Therefore, each point is assigned to the medoid that it is closest to.

The optimal number of clusters was determined through comparison of silhouette widths of real data to randomized data. In short, the data was subjected to PAM clustering for a given number of clusters. Randomized data was subjected to the same clustering 10 different times. Silhouette widths of the randomized tests were compared to that of the data and the optimal cluster number was determined as the number of clusters with the highest silhouette coefficient and widest range between experimental values and randomized controls with the aid of visual inspection to ensure optimal separation between clusters (Figure S2,S3). The silhouette width is a coefficient describing how well the clustering algorithm fits the data, with higher coefficients indicating a better fit.

#### Computational scaffold registration and integration

The 3D mapping data of the ICNS in the male and female F344 rats were curated by the SPARC data curation team. The curated data was then sent to the Auckland Bioengineering Institute (ABI) MAP-CORE branch for registering the 3D mapping data onto individual heart scaffolds. The registered data for two of the male F344 rat hearts standardize the 3D coordinates of the mapping data and to visualize the degree of similarity between ICNS distributions in common anatomical planes.

An anatomical scaffold mathematically defines the shape of an organ using a 3D *material* coordinate system. Within this coordinate framework many different aspects of tissue structure can be assembled including muscle fibre orientation, neural pathways, and the spatial distribution of ICNS data. Note that the term ‘material’ is used because these coordinates effectively identify the position of any material (tissue) particle, independent of its particular location in 3D space or how distorted it is. This material embedding provides a powerful reference frame to analyze and compare the pattern of individual ICNS distribution on one integrated scaffold.

The generic topology of the atria scaffold was algorithmically generated from a set of anatomical and mathematical parameters using an open-source Python library called Scaffoldmaker. The customized generic atria scaffold was fitted to each subject’s image contours using a least squares optimization. Specifically, the sum of the weighted distances between each segmentation data point on the contour and its projection onto the nearest element was minimized during the fitting process. This distance is a function of the scaffold parameters. The fitted scaffold is able to capture the spatial location and distribution of ICNS neurons and embed them locally into the elements as material coordinates. This material embedding of ICNS neurons stores a unique one-to-one mapping that can be used to transform them onto the corresponding generic scaffold elements.

