## Supplementary figures and images for "3D single cell scale anatomical map of sex-dependent variability of the rat intrinsic cardiac nervous system"

### Supplementary Movie 1

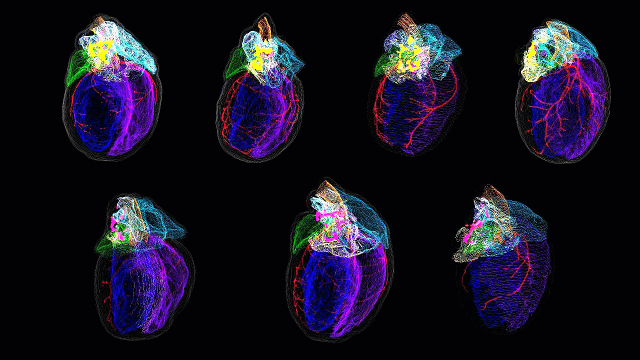
